# Inference based PICRUSt accuracy varies across sample types and functional categories

**DOI:** 10.1101/655746

**Authors:** Shan Sun, Roshonda B. Jones, Anthony A. Fodor

## Abstract

**Background:** Despite recent decreases in the cost of sequencing, shotgun metagenome sequencing remains more expensive compared with 16S rRNA amplicon sequencing. Methods have been developed to predict the functional profiles of microbial communities based on their taxonomic composition, and PICRUSt is the most widely used of these techniques. In this study, we evaluated the performance of PICRUSt by comparing the significance of the differential abundance of functional gene profiles predicted with PICRUSt to those from shotgun metagenome sequencing across different environments.

**Results:** We selected 7 datasets of human, non-human animal and environmental (soil) samples that have publicly available 16S rRNA and shotgun metagenome sequences. As we would expect based on previous literature, strong Spearman correlations were observed between gene compositions predicted with PICRUSt and measured with shotgun metagenome sequencing. However, these strong correlations were preserved even when the sample labels were shuffled. This suggests that simple correlation coefficient is a highly unreliable measure for the performance of algorithms like PICRUSt. As an alternative, we compared the performance of PICRUSt predicted genes to metagenome genes in inference models associated with metadata within each dataset. With this method, we found reasonable performance for human datasets, with PICRUSt performing better for inference on genes related to “house-keeping” functions. However, the performance of PICRUSt degraded sharply outside of human datasets when used for inference.

**Conclusion:** We conclude that the utility of PICRUSt for inference with the default database is likely limited outside of human samples and that development of tools for gene prediction specific to different non-human and environmental samples is warranted.

## Introduction

Recent advances in next generation sequencing are revolutionizing our understanding of complex microbial communities. Amplicon sequencing of marker genes provides information regarding the phylogenetic diversity and taxonomic composition of microorganisms present in the environment, while shotgun metagenome sequencing provides additional information on the relative abundance of functional genes. Although knowledge of taxonomy and functional genes of microorganisms are both important, functional genes are more directly related to pathways and therefore are essential for understanding the roles microorganisms play with regards to different physiological or ecological outcomes. However, the higher cost of metagenome sequencing hinders its application in studies consisting of a large number of samples, which are usually necessary in order to ensure adequate statistical power for detecting true differences [1]. Additionally, metagenome sequencing can also be very challenging for low biomass samples or samples that are dominated by non-microbial DNA [2, 3].

In order to address this problem, tools have been developed to predict microbial functional genes from their taxonomic compositions inferred from more cost-effective amplicon sequencing, including PICRUSt, Tax4Fun and FaproTax [4–6]. Among these tools, PICRUSt is the most widely used and has been applied in hundreds of projects on various environments, including human gut [7, 8], murine [9, 10], fish [11], coral [12], water [13], plant [14], bioreactor [15] and soil [16]. PICRUSt predicts the genes of organisms without sequenced genomes based on mapping their 16S rRNA genes to homologous taxa with fully sequenced genomes. The predictions of PICRUSt are therefore limited by currently available genomes, which are highly biased towards microorganisms associated with human health and biotechnology use [17].

In order to gauge the reliability of PICRUSt predictions in different environments and for different functional categories, we utilized human, non-human animal (gorilla, mouse and chicken) and environmental (soil) datasets that were sequenced for both 16S rRNA marker genes and shotgun metagenomes. We compared the predicted functional profiles from PICRUSt to the functional profiles measured with shotgun metagenome sequencing. We demonstrated that simple correlations such as Spearman correlation overstate the accuracy of PICRUSt by not taking into account the low variance of functional profiles generated from shotgun metagenome sequencing. As an alternative metric, we used PICRUSt results for inference with simple statistical models and found reasonable performance for human datasets, which presumably reflected the better reference information we currently have for human genomes, but a sharp decrease in performance for inference in non-human samples.

## Results

### Spearman correlation is not a reliable measurement for the prediction accuracy of gene contents

In order to test the accuracy of PICRUSt from a range of environments, we compared PICRUSt predictions to the results of shotgun metagenome sequencing on publicly available datasets for which both metagenome and 16S rRNA sequences were available (Table S1). As we would expect from previous literature [4], gene content estimation from PICRUSt was robustly correlated with gene contents from metagenome sequencing with Spearman correlations in the range of 0.62 to 0.84 (Fig. 1). For example, in one soil sample (Fig. 1a), there is a clear correlation between the relative abundance of each gene from PICRUSt and the relative abundance from metagenome sequencing (Spearman’s rho = 0.85). However, if the same comparison is made with the sample labels shuffled, the correlation that was observed is not substantially impacted (Spearman’s rho = 0.84) (Fig. 1b).

**Fig. 1.**
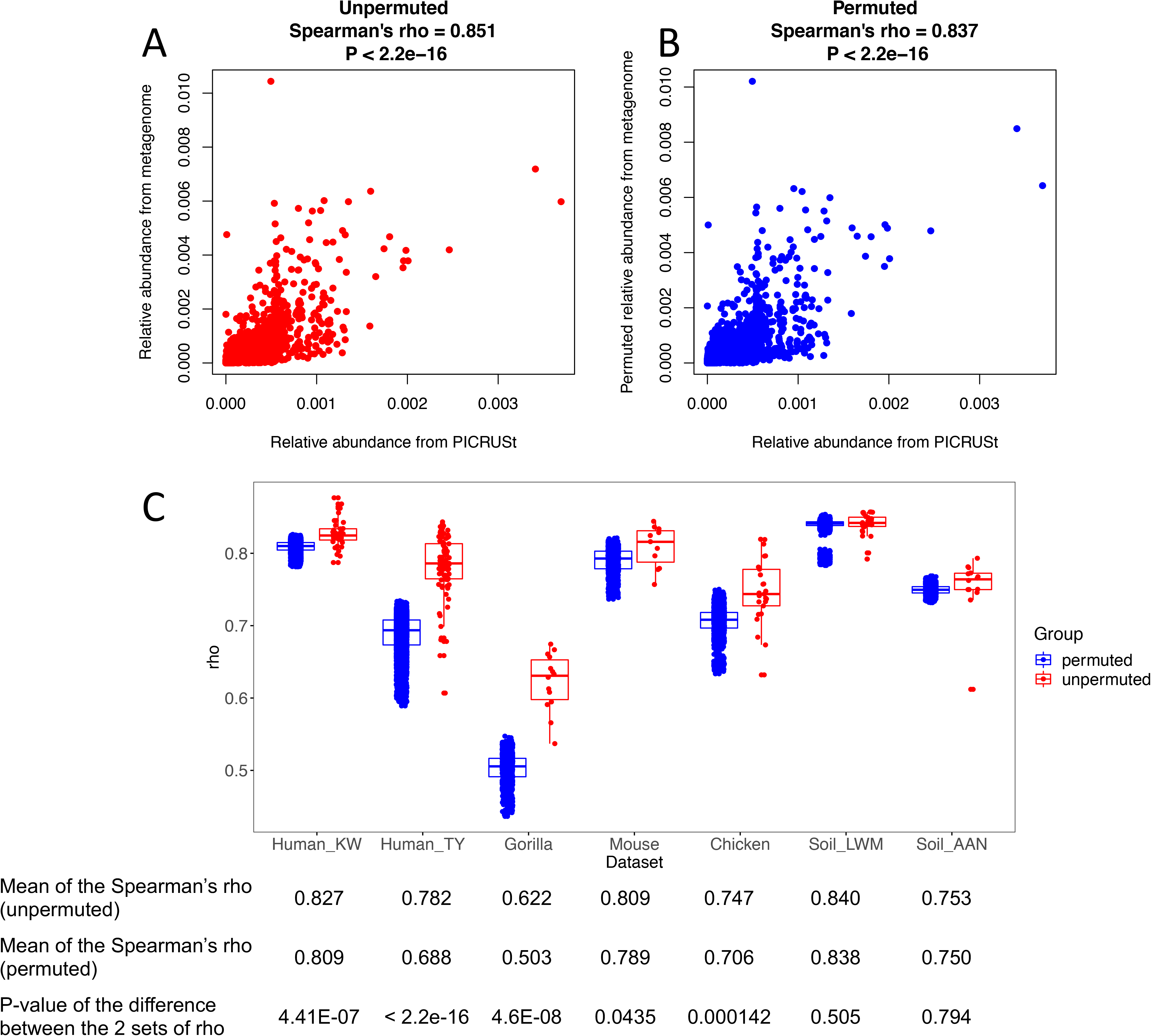
Spearman correlations between PICRUSt and shotgun metagenome sequencing in unpermuted and permuted datasets. **A** and **B:** An example showing the correlations between genes relative abundances estimated by PICRUSt and metagenome sequencing in a soil sample (sample BulkAG3 in soil_AAN dataset) for unpermuted (**A**) and permuted (**B)** sample labels. **C:** Comparison of Spearman’s rho between PICRUSt and metagenome sequencing in unpermuted (red) and permuted data (blue) in all datasets. In each of the 100 permutations, all sample labels were shuffled for every gene.

The likely explanation for this observation is that across environments, there is less variation between metagenome functional profiles of samples than their taxonomic profiles (Fig. 2), an observation that has been previously made for human samples in the Human Microbiome Project [18]. In the datasets we examined, despite sample labels being shuffled, the relative abundance of genes from PICRUSt estimates were highly correlated with the relative abundance of gene estimates from metagenome sequencing, with correlation coefficients always higher than 0.5. The PICRUSt predictions were often only marginally higher on the unpermuted data than those with sample labels shuffled, with perhaps the gorilla dataset as an exception (Fig. 1c). However, even in the gorilla samples, the difference between Spearman coefficients for permuted and unpermuted samples was only 0.12. For the 2 soil datasets, the Spearman coefficients for the unpermuted samples were not significantly different from those for the permuted samples (P = 0.508 and 0.794).

**Fig. 2.**
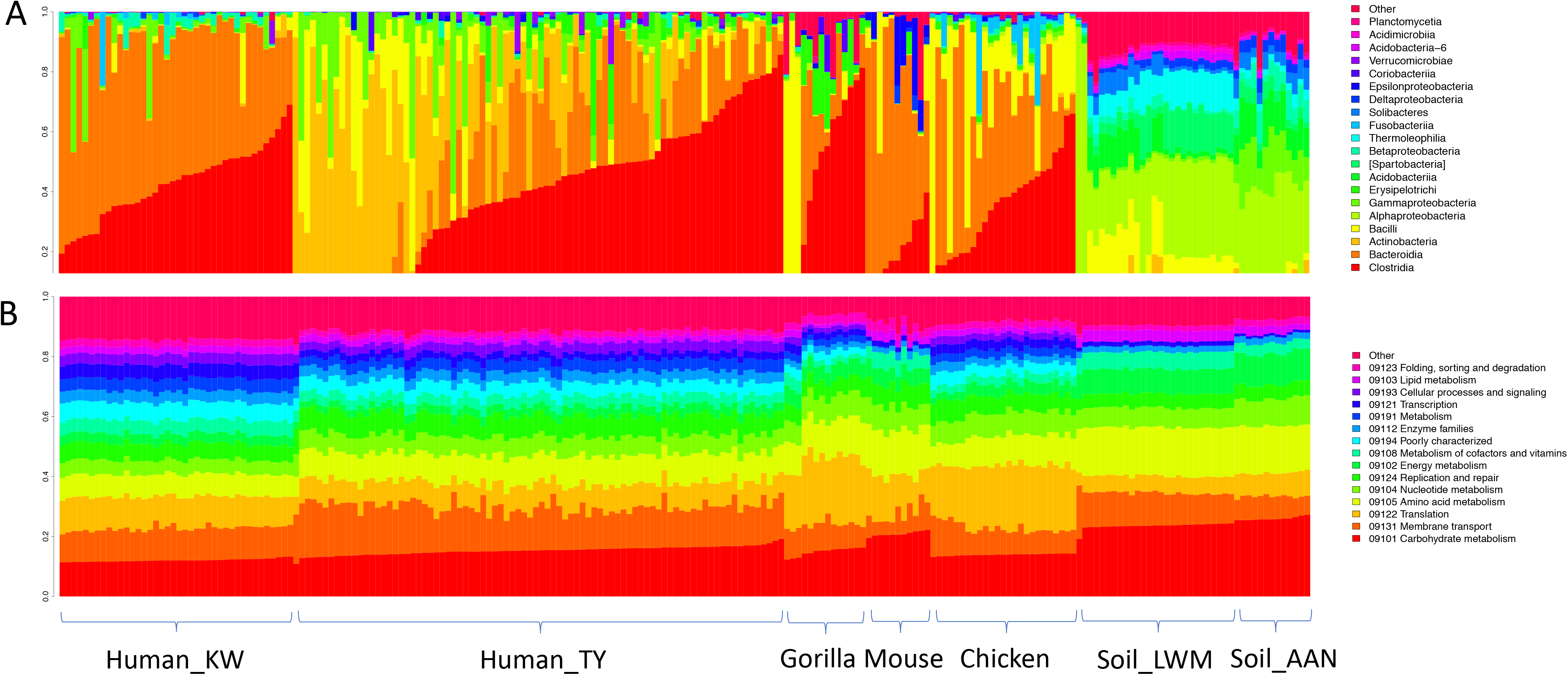
Taxonomic (**A)** and functional profiles (**B)** of the 7 datasets in our study. The taxonomic profiles were plotted at the class level and the functional profiles were plotted at the broadest functional category of the KEGG database for visualization.

### PICRUSt results for inference showed higher consistency with metagenome sequences in human samples than non-human samples

As an alternative evaluation to Spearman correlation, we compared how PICRUSt’s gene predictions performed in a t-test to genes detected with shotgun metagenome sequencing in each of our datasets. For this purpose, we formed a null hypothesis for each gene in each dataset that there is no difference in the mean of that gene’s distribution of relative abundance between the two groups in the dataset. For example, for each of the 5,574 genes detected by both PICRUSt and metagenome sequencing in the Human_KW dataset, we used a t-test to generate a P-value for the difference in gene composition between rural and urban samples. Across all the genes, there was a reasonable correlation (rho=0.49) of P-values from t-tests run on metagenome sequencing data and those on PICRUSt data (Fig. 3; top left panel). Unlike our results for Spearman correlation, this rho value is sensitive to sample permutation, as when we repeated this procedure on permuted data, the correlation between P-values generated from metagenome sequencing and those from PICRUSt approached zero (Fig. 4). We saw a similarly robust correlation (rho=0.554) for our Human_TY dataset evaluating a null hypothesis comparing US and non-US samples. However, when we extended this analysis to non-human datasets (using the null hypotheses for each study listed in Table S1), the inference produced by PICRUSt showed a much lower similarity to inference produced by metagenome sequencing (Fig. 3).

**Fig. 3.**
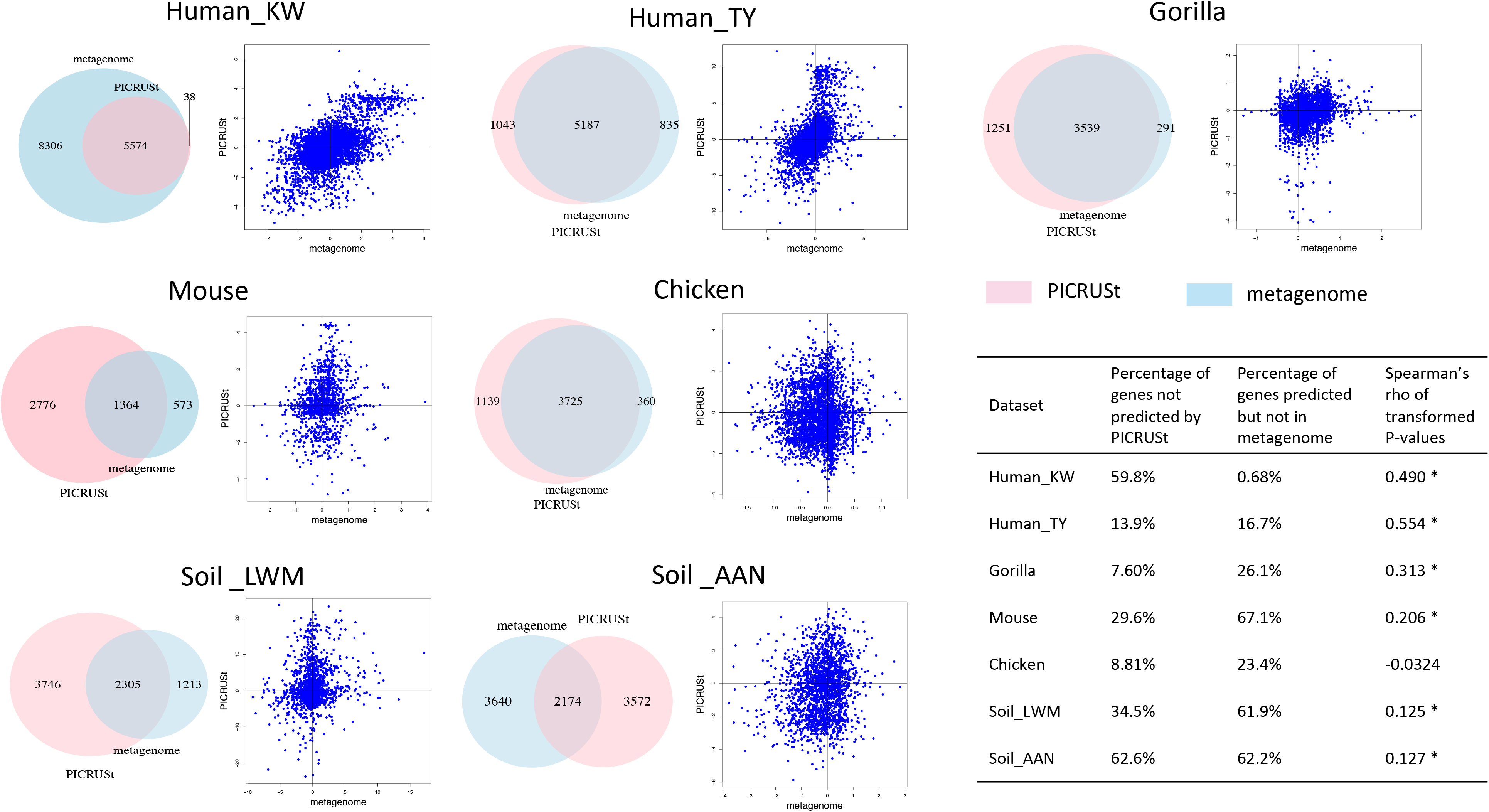
Comparison of inferences based on gene composition estimated with PICRUSt and metagenome sequencing in each of the 7 datasets. The Venn diagrams (left in each panel) show the number of genes detected by both PICRUSt and metagenome sequencing or by only one method, with those detected by PICRUSt in pink and those by metagenome sequencing in blue. The plots (right in each panel) show log-transformed P-values of the t-test evaluating the null hypothesis for each dataset (see methods and Table S1) from metagenome sequencing (x-axis) and PICRUSt (y-axis) for genes in common between the two methods. The sign of the log-transformed P-values reflect the direction of change (see methods). For example, in the Human_KW dataset, genes higher in urban subjects are in the upper-right hand quadrant and genes lower in urban are in the lower-right hand quadrant. The table (inset lower right corner) shows the percentage of genes called by only one method for each dataset and the coefficients of Spearman’s correlations between log-transformed P-values based on gene composition estimated by PICRUSt and metagenome sequencing.

**Fig. 4.**
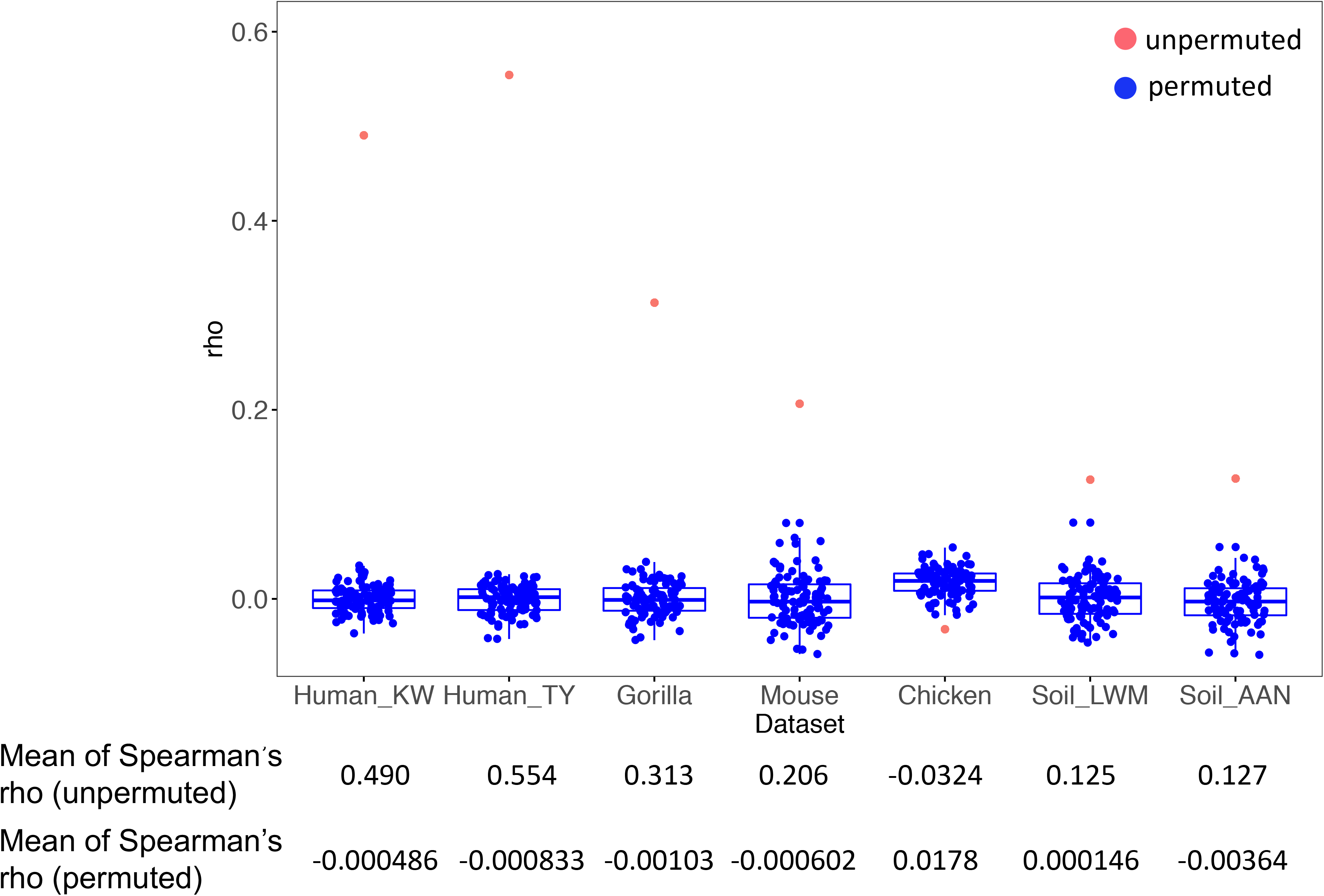
The results of inference methods in unpermuted and permuted datasets. The red points are the Spearman’s rho of log-transformed P-values from PICRUSt and unpermuted metagenome sequencing data for each dataset. The boxplots of blue points show the Spearman’s rho of log-tranformed P-values from PICRUSt and permuted metagenome sequencing data in 100 permutations. In each of the 100 permutations, all sample labels were shuffled for every gene.

In order to ensure that the differences in performance were not due to differences in sample sizes, we randomly sub-sampled each larger dataset (without replacement) to 10 samples (5 per group) and re-calculated the comparison of P-values between PICRUSt and metagenome sequencing (Fig. S1). Even at a smaller size, data from the human studies showed greater concordance than those from other environments. We conclude that the difference in sample sizes between datasets does not explain the variability of PICRUSt accuracy between different samples types in our study. Likewise, the effect sizes of the associations with metadata, measured as R^2^ in a PERMANOVA test, were not substantially higher in human samples (Table S1). It therefore also seems unlikely that effect size alone can explain the better concordance we observed between inference results from PICRUSt and metagenome sequencing for human samples.

In order to further investigate the consistency of PICRUSt and metagenome sequencing, we next asked how many genes were detected by both methods or by each method individually. For some datasets, such as the Human_KW dataset, PICRUSt failed to predict many genes that were detected by metagenome sequencing. For other datasets, such as the soil datasets, many genes that PICRUSt predicted were not detected in metagenome sequencing and there were also many genes seen in metagenome sequencing but not in PICRUSt (Fig. 3). For the chicken dataset with an average metagenome sequencing depth of 31 million reads/sample and the gorilla dataset of 27 million reads/sample, 23.4% and 26.1% of PICRUSt predicted genes could not be detected by metagenome sequencing (Fig. 3). In addition, the metagenome sequencing of the Human_KW dataset with an average sequencing depth of 10 million reads/sample detected 13,880 genes and PICRUSt missed 59.8% of them (Fig. 3).

### PICRUSt performs differently for different functional categories

We next investigated the discrepancy between PICRUSt and metagenome sequencing for inference in different functional categories (Fig. 5). When comparing the P-values from PICRUSt data to P-values from metagenome sequencing, there was a lack of significant correlations in the human gut samples for some categories including Signaling molecules and interaction, Immune system, Endocrine system, Cell growth and death, Nervous system, Digestive system, Environmental adaptation, Cardiovascular diseases, Immune diseases, Substance dependence, Circulatory system, Excretory system and Viral protein family. PICRUSt appeared to have a closer match to the metagenome sequencing data for categories such as Folding, sorting and degradation, Translation, Transcription, Replication and repair, Nucleotide metabolism, Glycan biosynthesis and metabolism, Drug resistance, Neurodegenerative diseases, Endocrine and metabolic diseases and Aging. For the genes only detected by one method, most of the genes missed by PICRUSt belong to Environmental Information Processing, Organismal Systems and Human Diseases, while those associated with Metabolism were more likely predicted (Fig. S2). Among the genes predicted by PICRUSt but not detected by metagenome sequencing, most of them belong to Cellular processing and signaling, Signaling molecules and interaction, Genetic information processing, Substance Dependence and Viral protein family (Fig. S3).

**Fig. 5.**
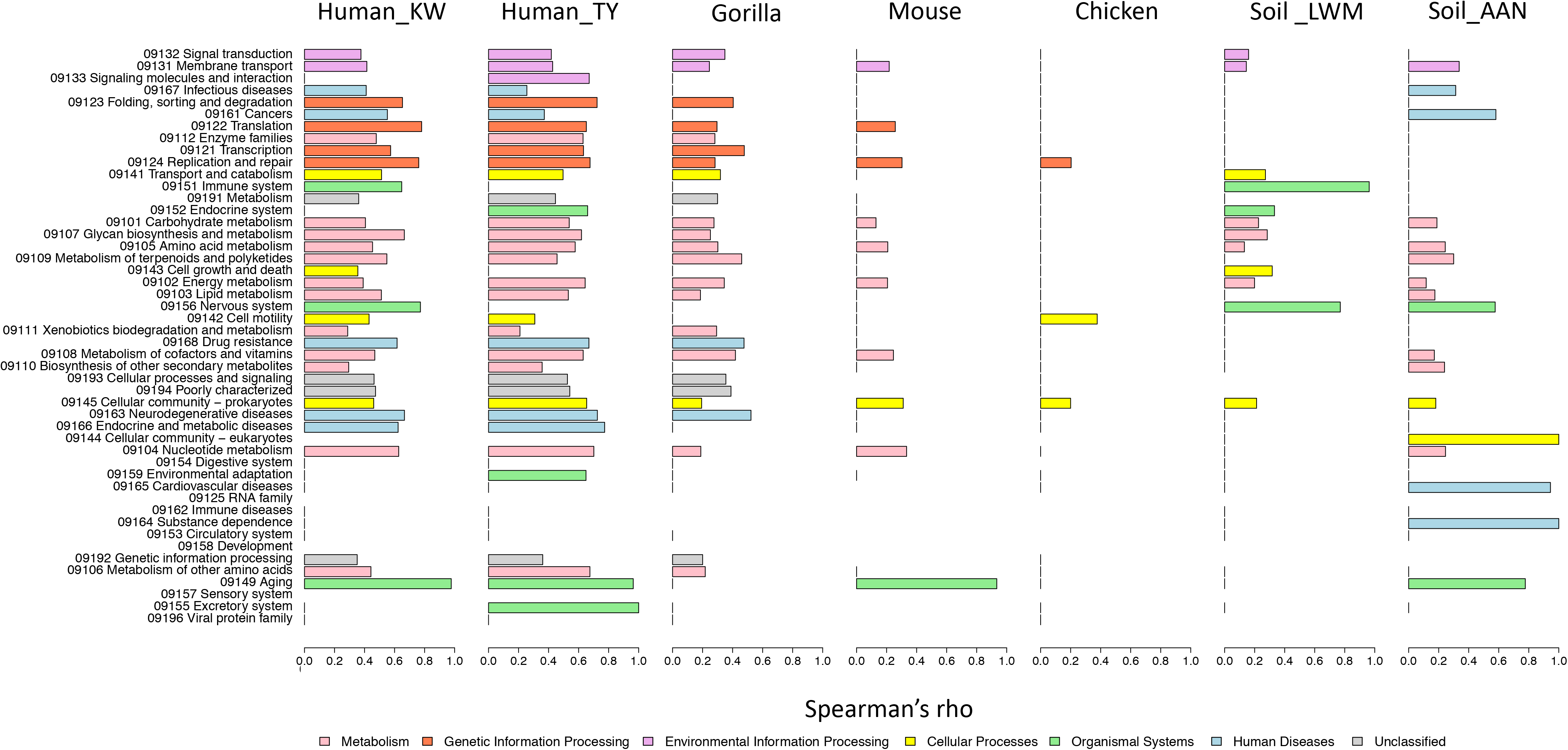
Spearman’s rho of the correlations between log-transformed P-values from PICRUSt and metagenome sequencing in 48 KEGG functional categories at the second hierarchy level with the bar colors indicating the functional categories at the first hierarchy level.

## Discussion

Microbial community functional profiles are typically of much lower variance compared to their taxonomic profiles likely because of the large proportions of “core” or “housekeeping” functions [18–20]. In this study, we showed that this lack of variance in functional profiles between samples leads to a strong correlation between functional profiles from metagenome sequencing and those estimated from references with PICRUSt, even when the sample labels are randomly shuffled (Fig. 1c). We argue that this result shows that metrics commonly used to measure gene prediction performance, such as Spearman correlation between gene composition estimated with PICRUSt and metagenome sequencing, do not give a satisfactory measure of overall accuracy. As an alternative, we evaluated the performance of PICRUSt at a community level based on inference from simple statistical models testing the association between genes and metadata. Unlike simple Spearman correlations, evaluation with inference methods are highly sensitive to shuffling sample labels (Fig. 4), which indicated that inference methods are much less affected by the relatively low variance of functional profiles. The inference-based approach also has the advantage of reflecting the common use of PICRUSt to reveal predicted functional profiles associated with different metadata categories [11, 12, 21–23]. Incorrect estimation of differential abundance could lead to false discovery of signature genes, and this concern motivated our approach to determine the reliability of PICRUSt produced inference in different systems.

In this study, we selected 7 datasets from different environments which include human, non-human animal and environmental (soil) samples. With inference methods, we found that PICRUSt and metagenome sequencing had more consistent assessment from human datasets than non-human animals or environmental datasets. It is likely that these differences reflected the bias of genome databases towards human-related microorganisms. However, PICRUSt still missed a large percentage of genes that were detected with metagenome sequencing in human samples, and an increase in metagenome sequencing depth could presumably increase the number of genes that are potentially not detected by PICRUSt (Fig. 3; Table S1). Likewise, PICRUSt sometimes predicted many genes not found in metagenome sequencing even in samples with presumably adequate sequencing depth of millions of reads per sample, which suggested that these additional genes are likely false predictions.

As a meta-analysis across multiple studies, there are systemic factors that may influence the results of this study, including different sample sizes, sequencing designs and effect sizes of associations with the metadata. We repeated our analysis on subsampled datasets that were rarified to the number of samples in the smallest dataset that we examined and observed a similar pattern of results with inference more consistent between PICRUSt and metagenome sequencing for human studies (Fig. S1). This result suggests that difference in sample size does not explain the better inference performance of PICRUSt for the human studies. While differences in effect size and experimental design are harder to control, the human studies did not have an obviously higher effect size than the non-human studies as measured with a PERMANOVA test (Table S1). It therefore also seems unlikely that differences in effect sizes of associations with the metadata can explain our results.

Our study also examined the performance of PICRUSt for different functional categories. This approach was motivated by the presumed bias in current genome databases toward culturable microorganisms [24]. We reasoned that the unculturable state of microorganisms could be caused by specific requirements for nutrients, temperature, pH, beneficial interactions with other microbes or extremely slow growth rates [25], which in turn could lead to bias in gene families in different microorganisms. Likewise, different microorganisms and genes also have different rates of horizontal gene transfer and the accuracy of gene content estimation may therefore vary depending on the type of the genes and microbial groups [26]. We found that PICRUSt performed best for “house-keeping” functions such as transcription and translation while the accuracy of functions related to environmental information processing was generally much lower (Fig. 5). Future algorithms for gene prediction could explicitly incorporate this performance variance into a confidence score that could give users estimated error rates for prediction of a given gene family.

Our analysis suggests that in order to better predict microbial functional profiles in certain environments, it will be of utility to develop tools specific to that environment. There have been encouraging examples in the literature of efforts to make environmental specific databases such as CowPI, a functional inference tool specific to the rumen microbiome, which had better estimates than PICRUSt when used for predicting functional profiles in the bovine environment [27]. We can look forward to similar future refinements in the next generation of these algorithms that will use appropriate reference databases for an environment and analyze individual functional categories to yield confidence scores for each prediction.

## Methods

The datasets used in this study include 2 human datasets (named as Human_KW [28] and Human_TY [29] in our study after the initials of their first authors), 1 gorilla [30], 1 mouse [31], 1 chicken [32] and 2 soil datasets Soil_LWM [33] and Soil_AAN [34]. Each dataset has publicly available 16S rRNA and metagenome sequences and is associated with a two-level categorical metadata. The Human_KW study compared urban and rural subjects in China, while US and non-US subjects were compared for the Human_TY study. In the gorilla study, the dry and wet seasons were compared while the mouse study compared community composition of two enterotypes. Lean and fat broiler chicken lines were compared for the chicken study. For the Soil_LWM study, Amazon dark earth and agricultural soil were compared, while forested and deforested soils were compared for the Soil_AAN study. Information regarding data locations, sequencing depth, sample sizes and effect sizes (measured as R^2^ in the PERMANOVA test with the function ‘adonis’ in the R package ‘vegan’) for each study are listed in Table S1.

The PICRUSt predictions of the 16S rRNA sequences in the datasets followed the developer’s instructions [4]. The authors’ metagenome analysis results were used when available [29, 31, 33, 34], otherwise the raw sequences were analyzed with humann2 following the developer’s instructions [35]. In each dataset, all PICRUSt-predicted gene families and pathways were compared to those from metagenome sequencing to determine their discrepancy in the number and types of genes revealed. For genes detected by both PICRUSt and metagenome sequencing, we used two sets of methods to evaluate their consistency. In a first set of methods, we analyzed the Spearman correlation between PICRUSt-predicted gene composition and those from metagenome sequencing. As a control, we permuted the sample labels 100 times and re-calculated Spearman correlation between PICRUSt estimates and metagenome sequencing estimates with sample labels shuffled.

In a second set of methods, we analyzed the consistency of PICRUSt and metagenome sequencing in the P-values they generated for null hypotheses of no association with metadata. For this purpose, P-values were produced with a t-test of the 2 distinguishable groups in each dataset (Table S1). P-values from the t-test were log10 transformed and multiplied by either 1 or −1 to include the direction of change as indicated below:

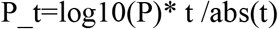

P_t is the transformed P-value, P is the P-value from t test, and t is the statistic of t-test. We then estimated the consistency of the log-transformed P-values from PICRUSt and metagenome sequencing with Spearman’s correlation. To determine whether this method is affected by the low variance of functional profiles, we permuted sample labels of the metagenome sequencing gene compositions 100 times and re-calculated the log-transformed P-values and their correlation with the PICRUSt results.

In order to correct for differences in sample size, each dataset was also subsampled to 5 samples per group to ensure that the different sample sizes of datasets were not unduly influencing our results. The PICRUSt predictions and metagenome sequencing were also compared in each of the 48 level 2 functional categories.

## Declarations

### Availability of data and material

The datasets analyzed in this study are publicly available with repositories and accession numbers listed in Table S1. R scripts used in this study are available at Github (https://github.com/ssun6/Inference_picrust). Additional requests and questions can be addressed to S.S.

### Competing interests

The authors declare that they have no competing interests.

