## SupplementaryFigures1-3 for "Inference based PICRUSt accuracy varies across sample types and functional categories"

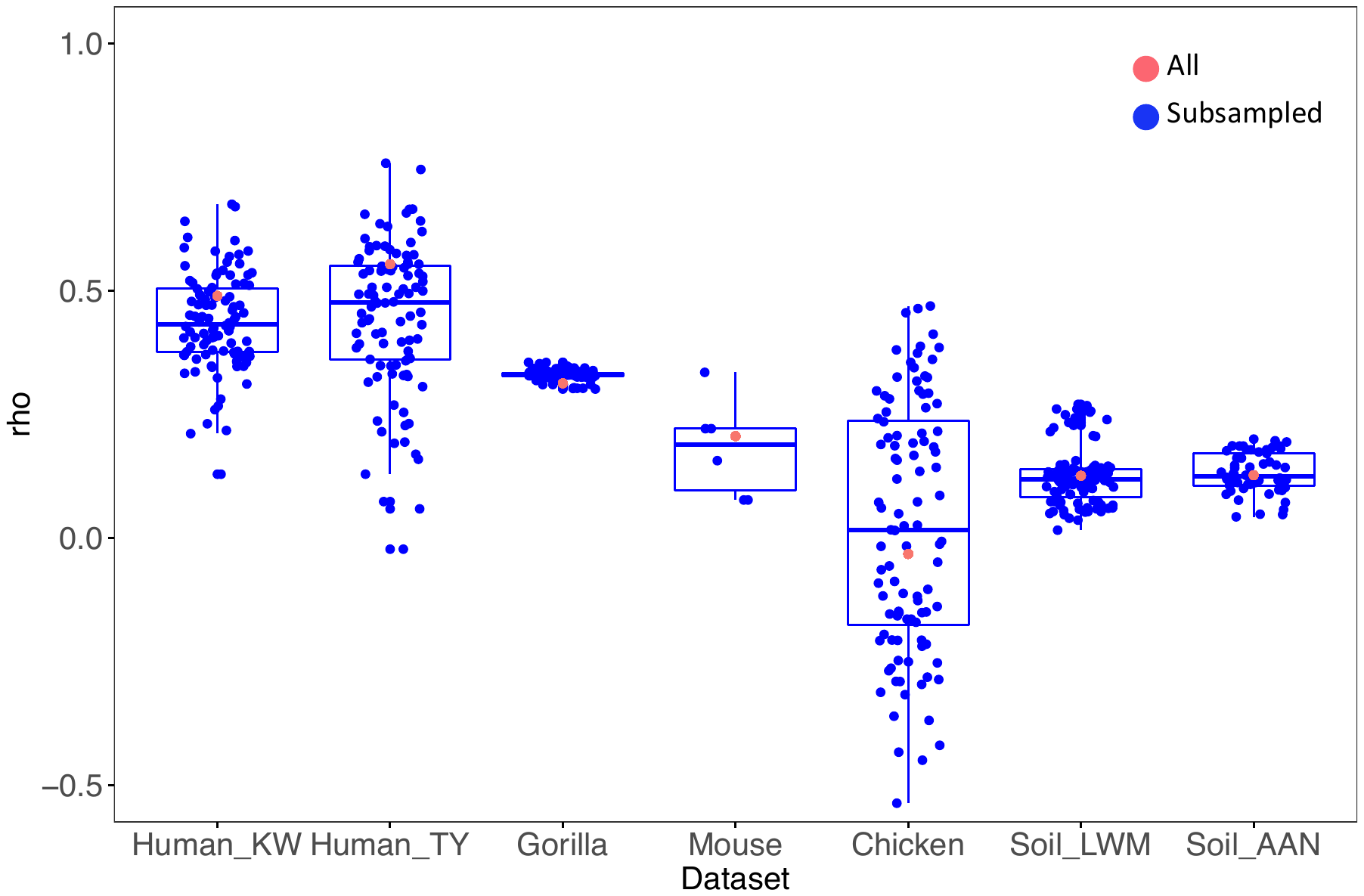

Fig. S1. The results of inference methods in complete and subsampled datasets. The red points are the Spearman’s rho of log-transformed P-values from PICRUSt and metagenome data in the complete datasets. The boxplots of blue points show the Spearman’s rho of transformed P-values from PICRUSt and metagenome data in subsampled datasets subsampled to 10 samples per study, with 5 for each group. The correlations were recalculated on 100 subsampled datasets when possible, otherwise it was recalculated on the maximum number of possible subsampling given the original sample sizes.

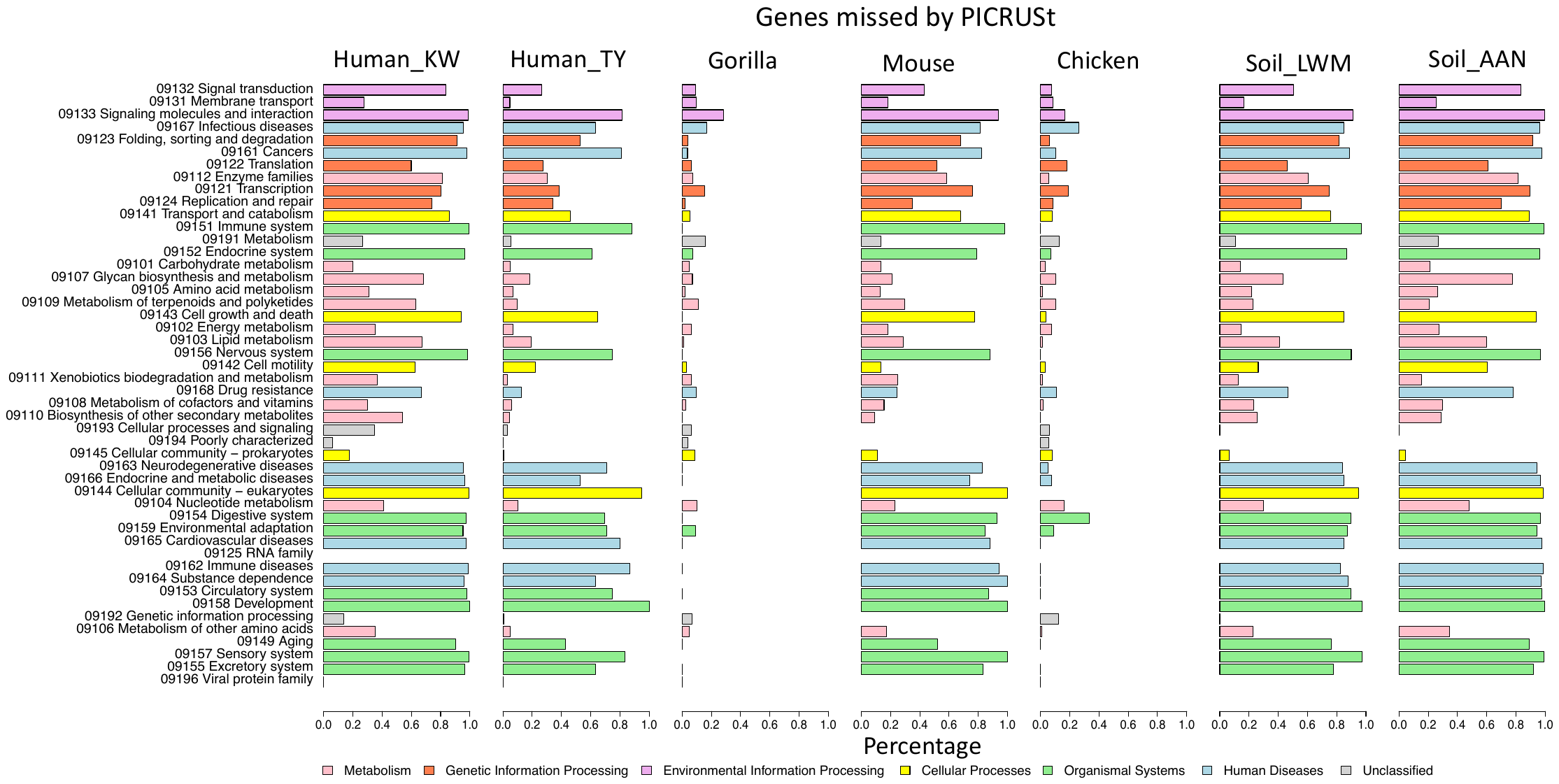

Fig. S2. The percentage of genes detected by shotgun metagenome sequencing but not by PICRUSt in 48 KEGG functional categories at the second hierarchy level with the bar colors indicate the functional categories at the first hierarchy level.

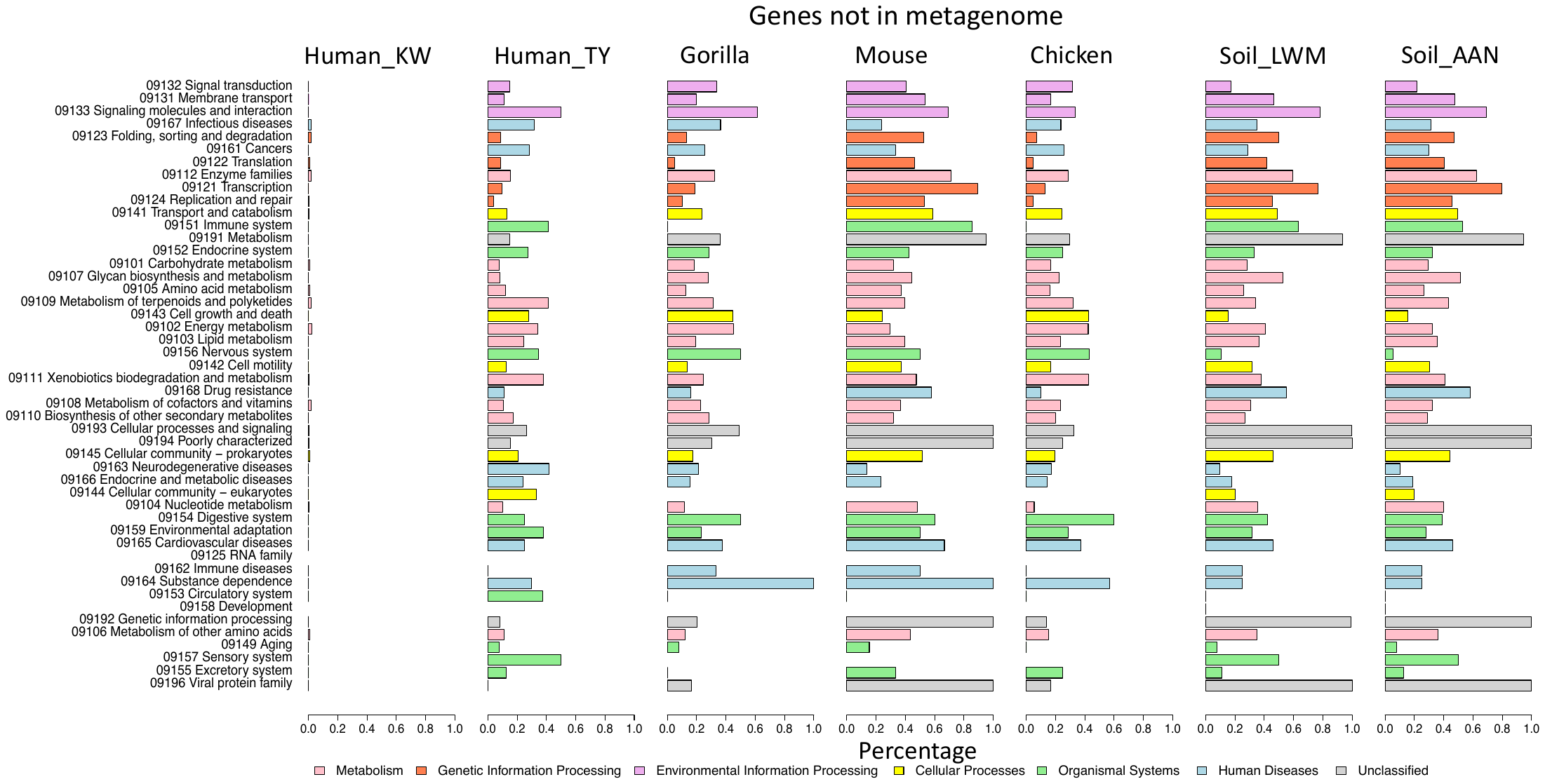

Fig. S3. The percentage of genes predicted by PICRUSt but not detected by shotgun metagenome sequencing in 48 KEGG functional categories at the second hierarchy level with the bar colors indicating the functional categories at the first hierarchy level.
